# Glymphatic clearance is enhanced during sleep

**DOI:** 10.1101/2024.08.24.609514

**Authors:** Erik Kroesbergen, Laurence V. Riesselmann, Ryszard S. Gomolka, Virginia Plá, Tina Esmail, Visti H. Stenmo, E. Rebeka Kovács, Elise S. Nielsen, Steven A. Goldman, Maiken Nedergaard, Pia Weikop, Yuki Mori

## Abstract

We here revisited the concept that glymphatic clearance is enhanced by sleep and anesthesia. Utilizing dynamic magnetic resonance imaging (MRI), single photon emission computed tomography (SPECT), and fluorescent fiber photometry, we report brain glymphatic clearance is enhanced by both sleep and anesthesia, and sharply suppressed by wakefulness. Another key finding was that less tracer enters the brains of awake animals and that brain clearance across different brain states can only be compared after adjusting for the injected tracer dose.

## Introduction

Over the past decade, a broad set of converging lines of evidence have supported the concept that clearance of brain waste proteins, including amyloid-β and tau, is increased during sleep and anesthesia. Direct injection of radiolabeled amyloid-β followed by brain removal and quantification of the residual radioactivity, revealed that sleep and anesthesia doubled amyloid-β clearance compared with awake mice ^1,2^. Elegant microdialysis studies then documented that the *in vivo* concentration of both amyloid-β and tau are higher in the awake and sleep deprived brain than during sleep ^3,4^. Neural synchrony promotes influx of CSF ^5,6^, and 3DISCO whole-body tissue clearing of mice showed that brain state markedly alters the distribution pattern of CSF tracers ^7^. In the human brain, sleep deprivation suppressed clearance of CSF contrast agents ^8^, while slow wave EEG activity during sleep was associated with the large scale movement of CSF into the 4^th^ ventricle ^9^. Also, PET imaging showed that amyloid-β accumulates in the human brain after one night of sleep deprivation ^10^. Despite these consistent findings, a recent study by Miao et al. challenged the established understanding that glymphatic activity peaks during sleep or anesthesia. Their report suggested that wakefulness, rather than sleep, enhances the spread of tracer within the brain ^11^. Here, we scrutinized the importance of brain state on glymphatic clearance by employing various imaging techniques to map tracer dispersion and glymphatic efflux in awake, sleeping, and anesthetized mice. Our use of 3D dynamic scanning revealed that awake mice exhibit significantly lower tracer delivery and dispersion within the brain compared to their sleeping and anesthetized counterparts. Crucially, we also observed that clearance to peripheral tissues is markedly reduced in awake mice. These findings reaffirm the existing body of literature, clearly documenting enhanced brain clearance during sleep and anesthesia.

## Results

An extensive adaptation and training protocol was developed to enable MRI imaging of awake mice, which were fitted with chronically implanted intrastriatal cannulas (**Fig. 1a**) for slow injection of a gadolinium (Gd)-based small contrast agent, gadobutrol (0.60 kDa) ^12^. Gd dispersion within the tissue was mapped during continuous dynamic contrast enhanced MRI (DCE-MRI) comparing awake and ketamine/xylazine (K/X) anesthetized mice. Gd spread to a greater extent from the site of injection and reached a larger volume of tissue in K/X anesthetized mice compared to awake mice (**Fig. 1b**). The Gd concentration was quantified in 4 concentric spheres located in increasing distances from the cannula, i.e. 0.0-0.5, 0.5-1.0, 1.0-1.5 and 1.5-2.0 mm. The Gd concentration reached a higher level in all 4 spheres in the K/X anesthetized mice (**Fig. 1c**), while the awake mice exhibited a delayed Gd arrival compared to K/X mice (**Fig. 1d**). The time-to-peak and peak concentrations exhibited significant differences, between the treatment groups, such that little to no contrast agent reached the outer spheres in the awake group (**Fig. 1e,f**). Together these findings show that much less Gd is delivered to the brain of awake compared to K/X anesthetized mice.

**Figure 1:**
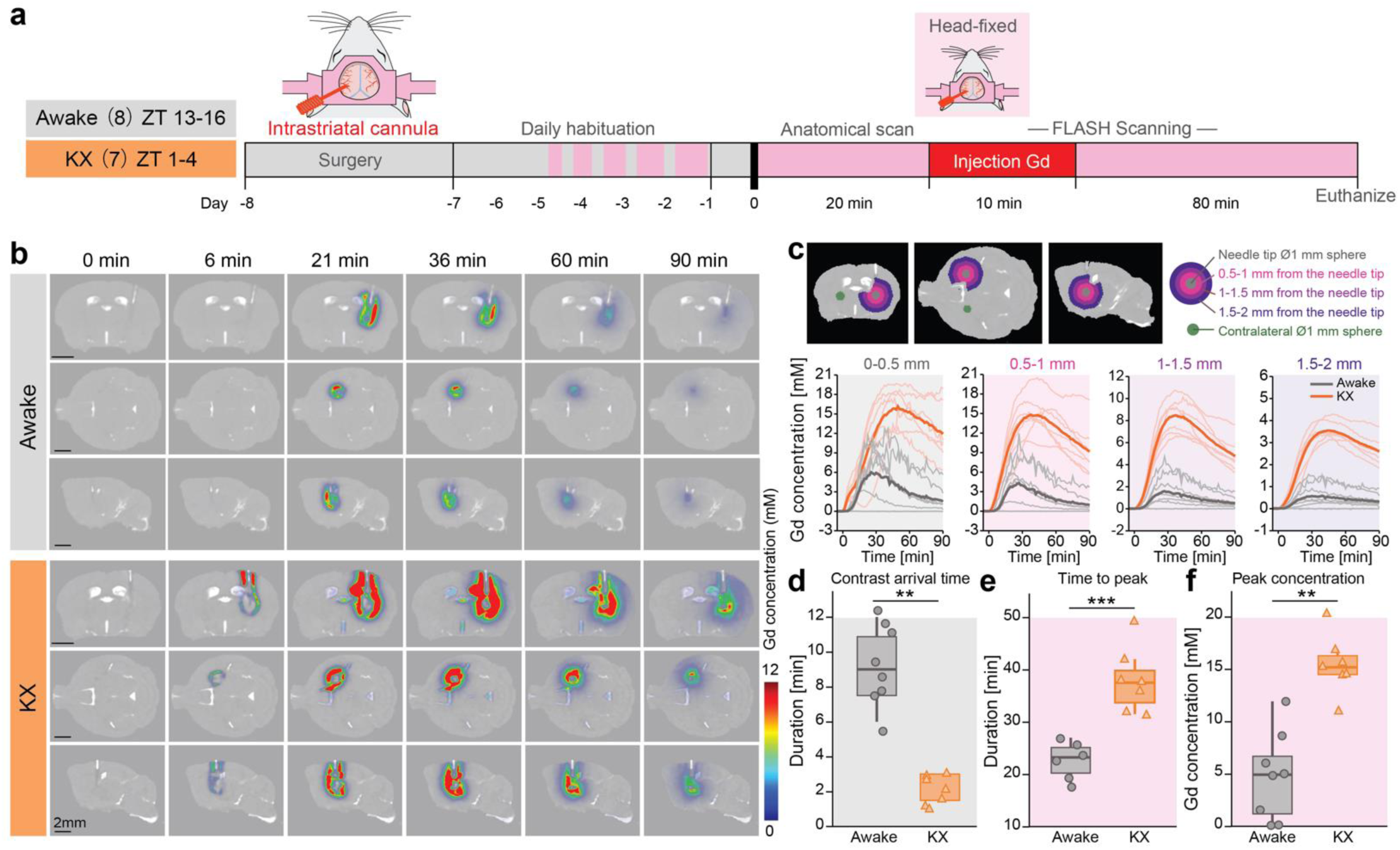
Anesthesia boosts the dispersion of intrastriate injected gadobutrol (Gd). **a,** Timeline representation of the DCE-MR imaging for awake and K/X anesthetized mice. Awake mice were studied at Zeitgeber time (ZT) 13-16, while K/X mice were studied at ZT 1-4. Gd (0.1 µl/min, 10 min) were injected in striatum at time 0. **b,** Representative 2D cross-sectional maps in 3 orthogonal planes showing the spatiotemporal distribution of Gd concentrations in awake and K/X anesthetized mice. The maps depict the concentrations at various times post-injection. **c, (upper)** The Gd concentrations were measured within concentric radius increments, 0-0.5 mm, 0.5-1.0 mm, 1.0-1.5 mm, and 1.5-2.0 mm, from the tip of the intrastriatal cannula. **(lower)** The concentration-time curves (CTCs) for Gd in the brain parenchyma, with solid lines representing group averages and dotted lines representing individual mice. Time points are marked from the start of the injection. **d,** Contrast arrival times, noted as the time when initial signal enhancement in the inner sphere (0.0-0.5 mm) were first detected after infusion pump started. If no tracer was detected within the 10 mins infusion period, the latency was recorded as “12 minutes”. **e,** Time-to-peak concentration in the sphere located 0.5-1.0 mmm from the cannula tip. **f,** Peak Gd concentrations, in the sphere located 0.5-1.0 mmm from the cannula tip. N = 8 (awake group), n = 7 (K/X-treated group). **-p<0.01, ***-p<0.001 using a non-parametric Mann-Whitney–U test.

We next used single photon emission computed tomography (SPECT) to detect spread of a radiotracer, ^99m^Tc-DTPA (0.49 kDa), infused into the striatum. Compared to MRI, SPECT is more sensitive and specific, registering only signal from the radiotracer. An initial CT/SPECT scan was obtained immediately after the 10 min radiotracer injection, to confirm delivery of the ^99m^Tc -DTPA (**Fig. 2a**). A second scan was then performed 3 hours post-infusion to assess the distribution of ^99m^Tc -DTPA within the brain (**Fig. 2b**). Between the two scans, the mice were left undisturbed in their cages between the two. This setup allowed for the inclusion of a natural sleep group in the experiment. EEG/EMG/video recordings were collected in parallel to capture the characteristic circadian regulation of the sleep-wake cycle (**Fig. S1**). The distribution volumes of ^99m^Tc -DTPA in sleeping and K/X-treated mice were 2.6- and 3.9-fold larger than those of the awake mice determined based on group-median volumes comparison (**Fig. 2c**). ^99m^Tc-DTPA dispersion was most extensive under K/X anesthesia, prominent in sleep, and virtually absent in awake mice. This variation highlights the impact of brain activity on tracer spread within the brain. Total ^99m^Tc-DTPA activity, measured in each mouse as a function the spherical distance from the infusion point, was highest in the K/X-treated mice, and lowest in the awake mice (**Fig. 2d**). In fact, the median activities recorded closest to the infusion point were over 15-fold higher in the K/X group and over 5-fold higher in the sleep group compared to the awake group. This difference in activity levels increased substantially with distance from the infusion point in the awake group and varied between 2.5- to 4-fold in the sleep group (**Fig. 2d**). Thus, the SPECT scans not only confirmed the MRI data using a highly sensitive alternative approach, but also extended the study by incorporating measurements from the natural sleep state.

**Figure 2:**
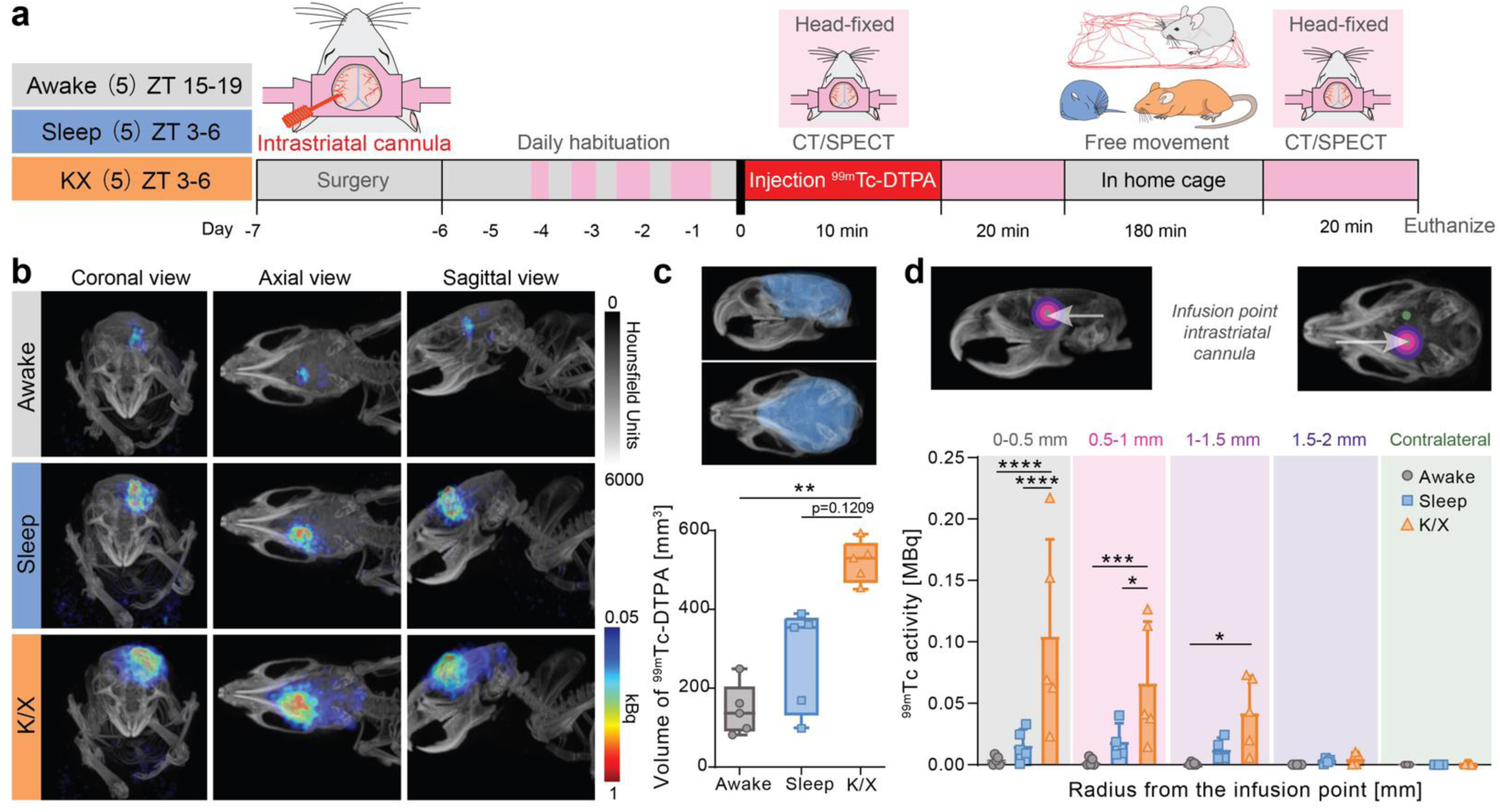
Sleep and anesthesia increase the tissue dispersion of ^99m^Tc activity. **a,** Timeline representation of the SPECT imaging experiment. ^99m^Tc-DTPA (0.1 µl/min, 10 min) was injected in striatum at time 0 and ^99m^Tc-DTPA distribution in brain was mapped after the mice spend 3 hrs in their cages. **b,** 3D maximum intensity projections of CT image from a representative mouse, along with volume reconstructions of mean SPECT images from all animals in the awake, sleep and K/X groups in coronal, axial, and sagittal views. **c,** Template brain parenchyma volume (blue) used for quantification of the ^99m^Tc-DTPA distribution volume **(upper)**, and whiskers-box plots for the total volume of activities present within the brain parenchyma of mice from the 3 groups **(lower)**. **d,** Bar plots of the total ^99m^Tc-DTPA activity present within the brain parenchyma of animals from the 3 groups, as measured within spheres located at 0-0.5, 0.5-1, 1-1.5, and 1.5-2 mm from the infusion tip. N = 5 in each group. *-p<0.05, **-p<0.01, ***-p<0.001, ****-p<0.0001 using non-parametric Kruskal-Wallis one-way ANOVA with Dunn’s post-hoc **(c)** or two-way ANOVA with Bonferroni’s post-hoc **(d)**.

The MRI and SPECT experiments focused solely on tracing dispersion within the brain, which does not equate to a measure of brain clearance. Brain clearance is defined by the outflow of substances to peripheral tissues. For instance, the distribution volume of a small tracer injected intrastriatally showed comparable distribution volumes in both AQP4 knockout and wildtype mice. However, clearance to peripheral tissues was reduced by 50% in the AQP4 knockout mice, highlighting a significant distinction between tracer distribution within the brain and its clearance to external tissues ^13,14^. We therefore developed a novel fluorescent approach to study tracer efflux to peripheral tissue in unrestrained awake, sleeping or anesthetized mice. Briefly, a small fluorescent tracer, FITC-dextran (3 kDa) was injected via a cannula in striatum, with concurrent assessment of tracer efflux to the periphery (blood) by means of an optic fiber implanted over the superior sagittal sinus (SSS), that had been placed 6 days earlier. Prior to tracer injection, the mice were habituated for 24 hrs to the experimental cage and further habituated for 1 hr to the cables connecting to the cannula and the optic fiber prior to tracer injection (**Fig. 3a, Fig. S2**). After the habituation period, the setup for data collection—including the pump, video recordings, and optical measurements—was activated remotely to avoid disturbing the mice. The awake group was studied during their active phase at ZT15-18, while experiments on the sleeping and anesthetized mice occurred during their inactive phase at ZT3-6 (**Fig. 3b**). Automatic analysis of the video recordings indicated that the awake mice were active for an average of 56.7 ± 5.4% (SEM) of the time over the 3-hrs duration of tracer circulation. This activity level was significantly higher compared to the naturally sleeping mice, who were active only 4 ± 0.6% of the time, and the anesthetized mice, who showed no movement (0%) (**Fig. 3b**). The FITC-dextran signal in the blood, indicative of brain clearance, showed a significant increase in the sleeping and anesthetized mice compared to awake ones during the 3-hrs *in vivo* recordings. Specifically, there was a 3.5-fold increase in the sleeping mice and a 2.4-fold increase in the anesthetized mice (**Fig. 3c**). Similarly, *ex vivo* analyses of blood collected at 3 hrs revealed a 2.8-fold increase in the sleeping group and a 2.1-fold increase in the anesthetized group (**Fig. 3d**). It is essential to note that we adjusted the FITC-dextran signal recorded from the SSS based on the injected tracer dose in each mouse to ensure accurate comparisons. At the conclusion of the 3 hrs recording period, the brains were collected, and the accumulated FITC-dextran in the optical nerves, trigeminal nerves, and superficial lymph nodes was imaged to assess efflux along the cranial nerves and lymphatic system. The results showed a notable increase in the FITC-dextran signal: sleeping mice exhibited a 2.5-fold increase, and anesthetized mice a 4.1-fold increase, compared to awake mice (**Fig. 3e**). Immunohistochemical analysis of the immersion-fixed brains indicated a non-significant trend toward mild astrogliosis (GFAP) and microgliosis (Iba1) in the area surrounding the cannula, with no significant differences observed between the awake, sleeping, and K/X treated groups. Additionally, neuronal density, as detected by NeuN labeling, was not reduced around the cannula site (**Fig. S3**).

**Figure 3:**
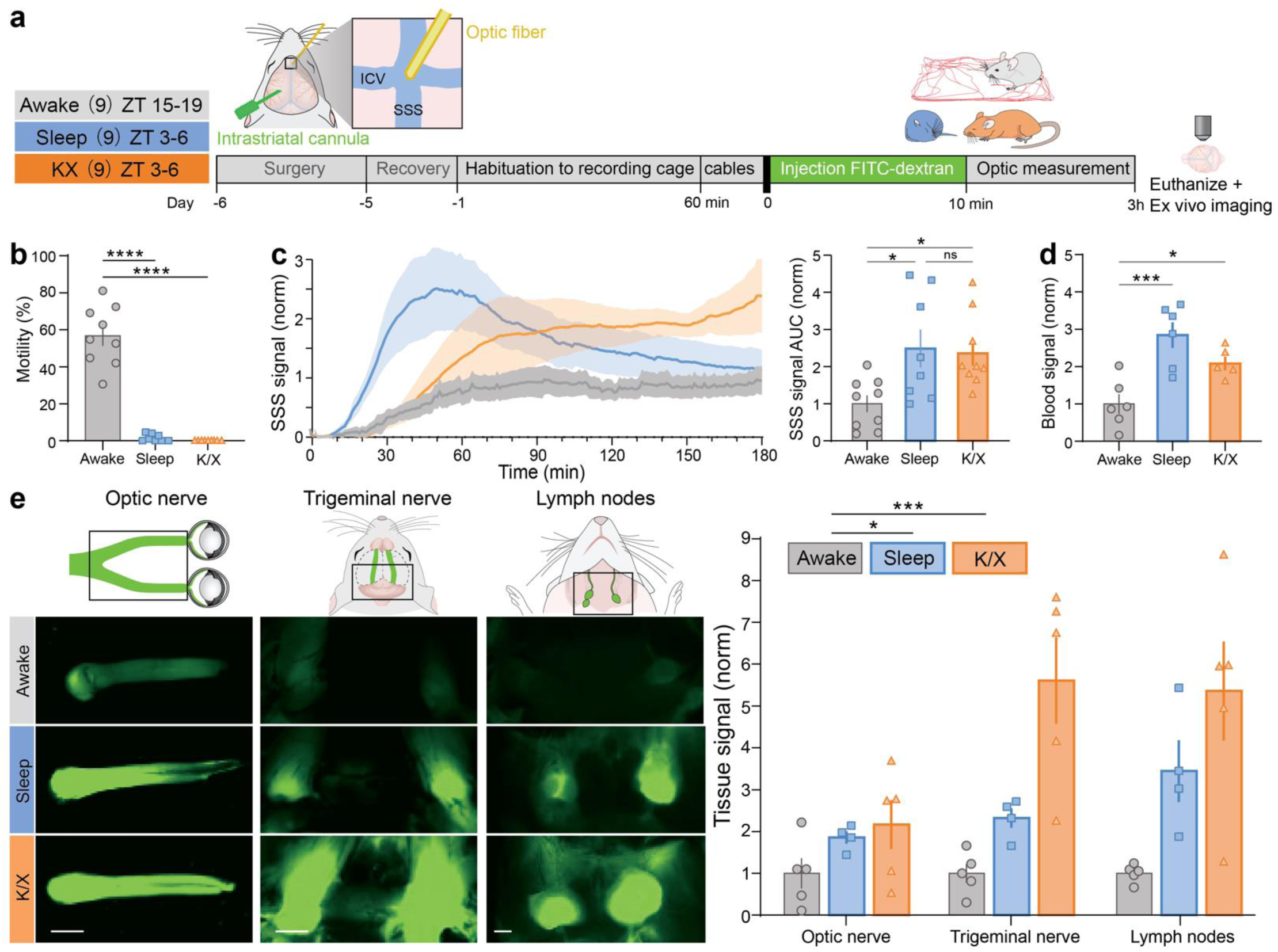
Clearance of intrastriate injected FITC-dextran is enhanced during sleep and anesthesia. **a,** Timeline representation of the fiber photometry experiment in awake, sleeping and anesthetized mice. FITC-dextran (3 kDa, 0.1 µl/min, 10 min) was injected in striatum at time 0. An optic fiber was placed over the confluency of the superior sagittal sinus (SSS) and the inferior cerebral vein (ICV) to detect FITC-dextran efflux (SSS signal) to the blood compartment. **b,** Quantification of the motility of the mice during the 3 hrs of tracer circulation. **c,** 3 hrs fluorescence signal detected by the optic fiber placed over the superior sagittal sinus (SSS) following intrastriate tracer injection after normalization for the injected volume **(left)**, and fluorescent signal under the curve of the *in vivo* recordings **(right)**. **d,** *Ex vivo* measurements of the fluorescence signal of blood samples in a subgroup of the mice. **e,** Representative images, and quantification of tracer efflux along the optic nerve, trigeminal nerve, and superficial cervical lymph nodes in a subgroup of the mice. N = 9 in each group. All data are represented as mean ± SEM, *-p<0.05, **-p<0.01, ***-p<0.001, ****-p<0.0001, using one-way ANOVA **(b-d)** or two-way ANOVA **(e)** with Bonferroni’s post-hoc.

Thus, three alternative measures of brain tracer efflux documented that clearance to the periphery is enhanced during sleep and K/X anesthesia and suppressed during wakefulness. This evidence collectively supports the notion that the brain’s waste clearance mechanisms are more active during states of unconsciousness.

## Discussion

This study confirmed that both natural sleep and ketamine/xylazine (K/X) anesthesia significantly enhance the clearance of small molecular weight tracers from the brain to peripheral tissues, with an increase of more than 2-fold compared to wakefulness. Additionally, it was found that wakefulness is associated with the suppression of tracer efflux, highlighting the distinct differences in brain clearance mechanisms across different states of consciousness ^1^. This conclusion is consistent with the extensive literature indicating that sleep and anesthetic regimens characterized by high delta power and low heart rate are associated with increased brain clearance ^15,16^. The findings emphasize the critical role of brain activity states in regulating the brain’s ability to remove waste, potentially offering insights into optimizing conditions for brain health and recovery. The contrasting conclusion recently reached by Miao et al. was based on two key pieces of evidence. First, fiber optic recordings from the prefrontal cortex demonstrated that a small fluorescent tracer injected into the caudate-putamen arrived later and with a lower peak amplitude in awake mice compared to those that were sleeping or anesthetized. The second line of evidence was derived from quantifying the tracer signal in coronal brain sections prepared 3 or 5 hrs post-injection. These sections showed that brains from awake mice contained much less tracer than those from sleeping or anesthetized mice. Based on the assumption that an equal amount of tracer was administered across the awake, sleeping, and anesthetized groups, the authors concluded that tracer clearance was enhanced in awake mice.

We here employed real-time imaging techniques to monitor tracer injection, dispersion, and clearance from the brain *in vivo*, utilizing several methods including dynamic contrast-enhanced MRI (DCE-MRI), radiotracer-based SPECT, and fluorescent fiber photometry. Dynamic 3D MRI results indicated that less tracer entered and dispersed within the brains of awake mice compared to those anesthetized with K/X (**Fig. 1**). High-sensitivity SPECT imaging further confirmed that the volume of injected radiotracer was smaller and exhibited poor dispersion in the brains of awake mice (**Fig. 2**).

The SPECT study was designed to allow tracer circulation while the mice remained unconfined in their cages, i.e., not subjected to head fixation, which enabled the inclusion of a group of naturally sleeping mice. This SPECT analysis revealed that wakefulness significantly suppresses both tracer injection and its dispersion within the brain compared with both sleep and K/X anesthesia.

Based on numerical analysis and 3D visualizations of the injected tracers via both MRI and SPECT, we conclude that the findings reported by Miao et al. likely stem from a reduced amount of tracer being administered to awake mice compared to those that were sleeping or anesthetized (**Fig. 1 and 2**). Focusing upon tracer dispersion in a single region using fiber optic (pinhole) techniques and limited the tracer quantification to only 3 and 5 hours post-injection, may obscure the effects of a reduce tracer injection volume in awake mice. Consequently, this approach could mistakenly suggest enhanced tracer clearance from the brain during wakefulness.

The analysis presented here incorporates a novel method for assessing tracer clearance to peripheral tissues in freely behaving mice. Detection of efflux of the intrastriate injected tracer in blood is based on the principle that ‘*all roads lead to Rome*’ – that all solutes cleared via the lymphatic system eventually drain into the bloodstream ^17^. By affixing an optic fiber atop the superior sagittal sinus, we enabled continuous monitoring of FITC-dextran export from the brain to the blood in awake and freely moving, naturally sleeping, and anesthetized mice. Our findings show that a small fluorescent tracer is cleared more extensively from the brains of sleeping and anesthetized mice compared to those that are awake (**Fig. 3**). Importantly, we ensured that the FITC-dextran signal in the blood was normalized to the volume of the tracer injected intrastriatally in each mouse. This careful normalization allows for accurate comparisons of tracer clearance across different physiological states. Based on multiple alternative imaging approaches, we conclude that brain clearance - defined as export of small solutes to peripheral tissues - is enhanced by natural sleep and K/X anesthesia.

---

**Figure S1:**
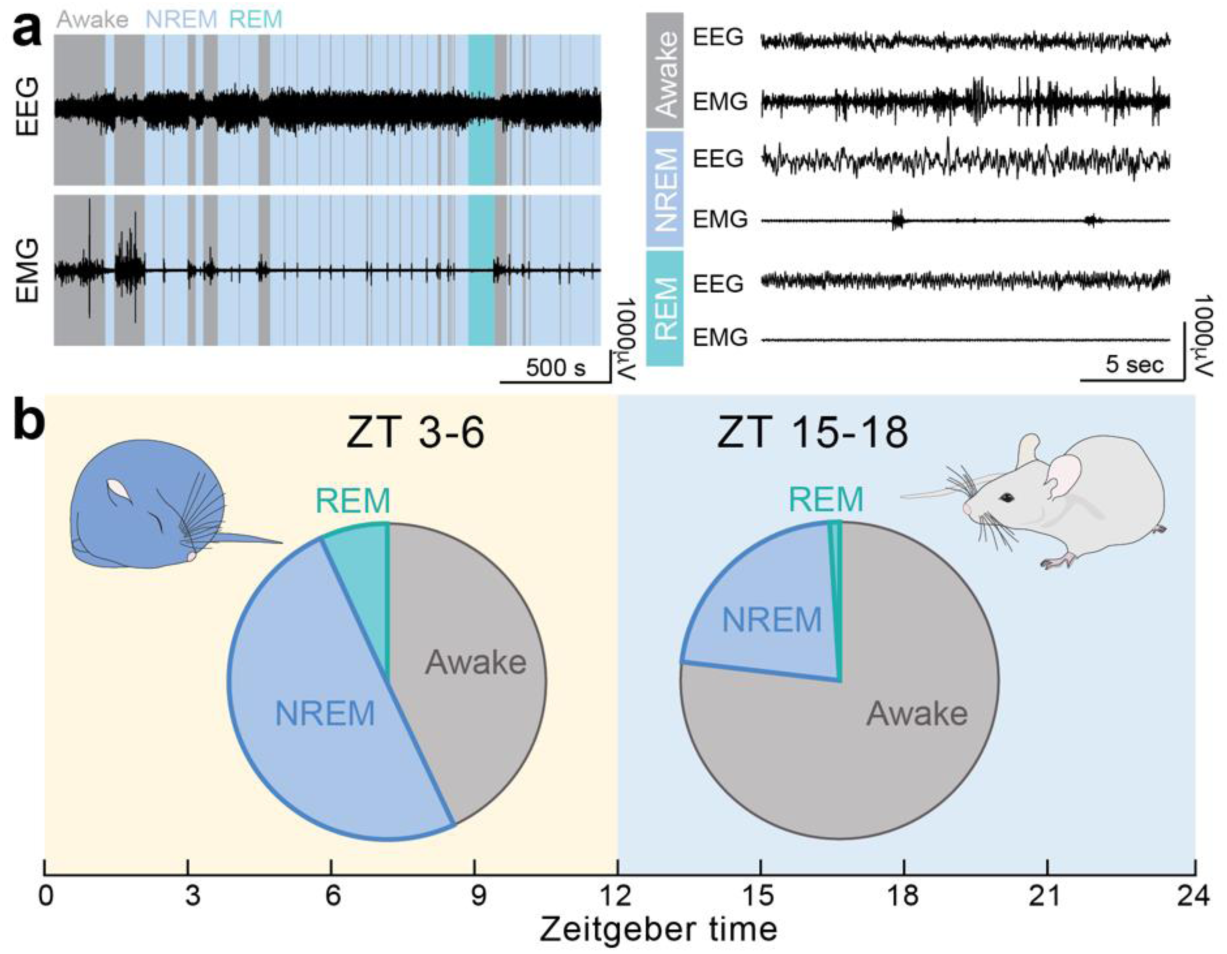
The circadian rhythm regulates the sleep-wake cycle. **a,** Representative recordings showing EEG and EMG activity during natural transitions between different brain states during the inactive phase **(left)**, and representative recordings showing typical EEG and EMG activity during wakefulness, NREM and REM sleep **(right)**. **b,** Percentage of time spent awake or in NREM and REM sleep at ZT 3-6 and 15-18. Hypnograms were generated from EEG/EMG traces scored in 1-second epochs. Five mice per group were analyzed, as previously described ^18^.

**Figure S2.**
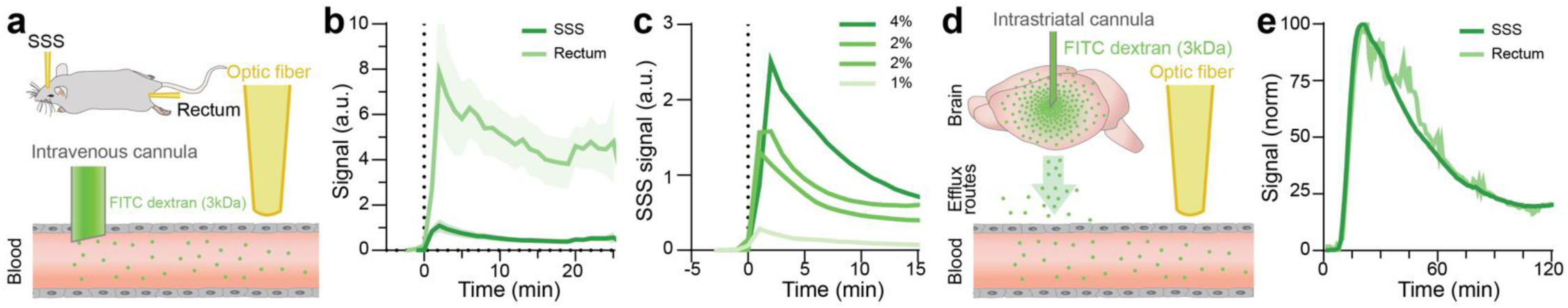
Validation of fiber photometry clearance assay. **a,** Schematic overview of fluorescent measurement at the superior sagittal sinus (SSS) and rectum after IV tracer injection. **b,** Comparison between FITC-dextran signal detected at the SSS and rectum in K/X anesthetized mice after injection of 1 µl 2% 3kD FITC-dextran IV (n = 7). **c,** Representative examples of dose-dependent fluorescent signal at the SSS in anesthetized mice after 1 µl of 1, 2 or 4% 3kD FITC-dextran IV injection. **d,** Schematic overview of fluorescent measurement at the SSS and rectum after intrastriate tracer injection. **e,** Representative example of normalized comparison between the FITC-dextran signal at the SSS and rectum in anesthetized mice after 2 µl of 10% 3kD FITC-dextran intrastriate tracer injection.

**Figure S3:**
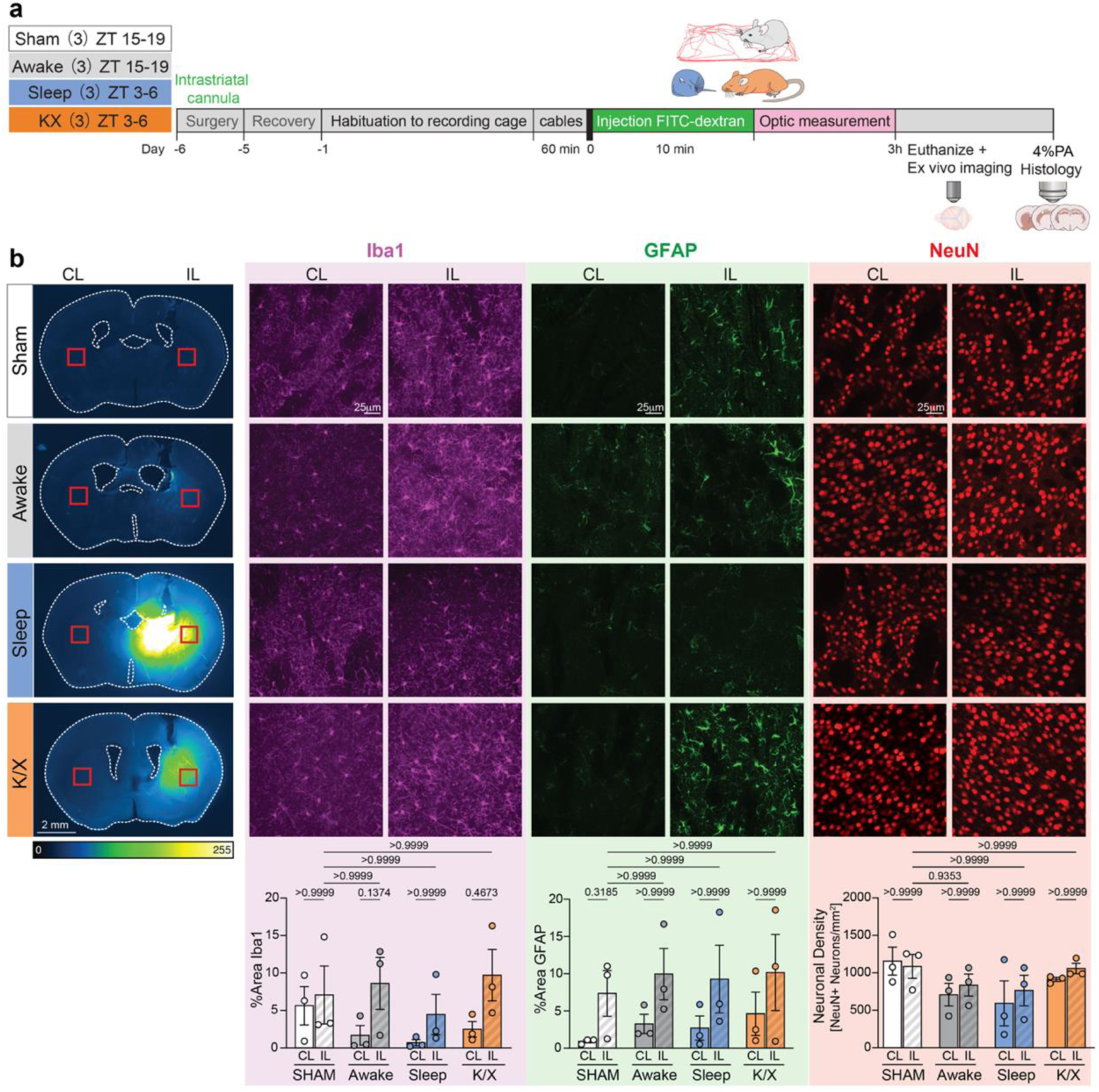
Impact of fiberoptic implant on reactive gliosis and neuronal density. **a,** Timescale of the performed experiments. In short, brains for histology were collected at the end of the fiberoptics experiments and the brains harvested 7 days after cannula implant (striatum, ipsilateral, IL), compared to an equivalent area in the contralateral region (striatum, CL). Animals receiving cannula implants, but no tracer were used as controls (Sham). **b,** Representative macroscopic images of complete brain slices are shown at the injection site, where the approximate location of the area used for reactivity evaluation is indicated (red squares) for each experimental group (Sham, Awake, Sleep and K/X). LUT is used to visualize the differences in tracer content. Representative high magnification confocal Z-stack projections are shown for both areas (CL and IL) where microglial and astroglial reactivity (Iba1, magenta and GFAP, green) was quantified. To evaluate the effect of cannular implantation on neuronal viability, neuronal density was measured as NeuN^+^ cells/mm^2^ (red, no significant differences). N = 3 in each group. Bar graphs summarize the data as mean ± SEM, with each individual dot representing one mouse. Non-parametric one-way ANOVA for matched samples (Friedman test), using Dunn’s multiple comparisons test, was employed for the statistical analysis. Scale bar is indicated on the images.

## Materials and Methods

### Animals

C57BL/6J mice of both sexes (Janvier Labs, Le Genest Saint-Isle, France), aged 9 to 14 weeks, were used for the experiments. All animals were group-housed (up to 5 mice per cage) prior to surgery and single-housed thereafter, with ad libitum access to food and water. The vivarium was maintained at a constant temperature of 22°C and controlled humidity. Mice studied during sleep or anesthesia were housed under a normal 12/12-hour light/dark cycle, while awake mice were housed in a room with an inverse 12/12-hour light cycle. All experiments were approved by the Animal Experiments Council under the Danish Ministry of Environment and Food (license numbers: 2020-15-0201-00559 and 2022-15-0201-01270). All procedures were conducted in accordance with the European directive 2010/63/EU.

### EEG/EMG/video and motility scoring

EEG and EMG recordings at Zeitgeber times (ZT) 3-6 and 15-18 were compared in a subgroup of mice. Low-impedance stainless steel screws (NeuroTek, 0.8 mm) were carefully inserted into two burr holes located above the frontal cortex and cerebellum (reference electrode) and secured to the skull with dental cement (SuperBond). Two silver wires (W3 Wire International) were inserted into the trapezius muscle to serve as EMG electrodes. The electrodes were implanted more than 2 weeks prior to recordings.

Mice were placed in a recording chamber (ViewPoint Behavior Technology) and connected to the EEG and EMG electrodes via cables linked to a commutator (Plastics One, Bilaney, SL12C), allowing free movement. The mice were acclimated to the recording chamber for at least 1 day before the experiments. Recordings were conducted for 3 hours, matching the duration and circadian timing of the fiber photometry recordings.

EEG and EMG signals were amplified (National Instruments, 16-channel AC amplifier, model 3500) and filtered (EEG: high-pass at 1 Hz and low-pass at 100 Hz; EMG: high-pass at 10 Hz and low-pass at 100 Hz). A 50 Hz notch filter was used to minimize power line noise. Signals were digitized using a Multifunction I/O DAQ device (National Instruments, USB-6343) at a sampling rate of 512 Hz. Continuous video recordings were captured using an infrared camera (FLIR Systems) and later used to aid in scoring vigilance states. EEG/EMG traces were scored in 1-second windows, with video recordings assisting the analysis. For SPECT/CT and fiber photometry experiments, motility-based sleep scoring was employed to minimize the need for implants that could interfere with brain fluid transport ^19^. Videos of cage activity were analyzed automatically for any movement of mice within their cage, based on pixel change (Ethovision).

### Surgeries: Headplating, cannula and fiber photometry implantation

General anesthesia was induced with 4% isoflurane and maintained at 1-3% isoflurane throughout all surgical procedures. Once the mice were unresponsive, they were placed in a stereotactic frame, and lidocaine (0.2 mg/ml) was administered locally at the surgical site. For additional analgesia, carprofen (5 mg/kg) and buprenorphine (0.05 mg/kg) were given subcutaneously. Intrastriatal cannulations were performed as described by Plá et al. ^13^.

In brief, after exposing the skull, a burr hole was drilled (M/L: 2.5, A/P: -0.3 relative to bregma) using a dental drill (Tech2000, RAM Digital Microtorque). A nylon cannula (0.24 mm inner diameter, Bilaney Consultants, UK) was then inserted to a depth of -3.2 mm. For mice enrolled in the DCE-MRI and SPECT/CT studies, a custom-made headplate was attached to the cranium immediately after cannula implantation.

For the fiber photometry clearance assay, a CMA/7 guide cannula with a dummy (CMA/Microdialysis, Stockholm, Sweden) was inserted more posteriorly into the striatum (M/L: 2.0, A/P: -0.6, D/V: -3.2 mm relative to bregma) to improve separation from the optical fiber. A ø200 µm, 1.5 mm optic cannula (Thorlabs) was positioned over the crossing of the superior sagittal sinus and the cerebral inferior vein, secured with dental cement. At this location, the venous sinus has a diameter of approximately 400-500 µm, compared to the 200 µm tip of the optic fiber.

Initial weights were recorded for all mice, and pain management was provided for 3 days post-surgery. After the procedures, mice were monitored closely, with daily weight measurements logged. Any mouse that did not recover well was excluded from the experiments.

### Adaptation and training protocols of headplated mice

**DCE-MRI:** After two days of post-surgery recovery, mice were trained for the MRI for five consecutive days with a 24-hour break before imaging. To account for the MRI environment, the habituation sessions started with 10 minutes of restraint on the first day and increased daily to reach 60 minutes on the fifth day using a mock MRI bed and restrain system ^12^. On the day of the experiment, the mice were briefly anesthetized with isoflurane, wrapped in gauze, and the headplate secured to the MRI mouse holder. **SPECT/CT:** Mice were progressively trained in the scanning bed, starting with 15-30 second sessions on Day 3, increasing to 5 minutes on Day 4, 15 minutes on Day 5, and 20 minutes on

Day 6. The training protocol was lighter since the maximal duration of restraining was shorter than used for the DCE-MRI studies. **Fiber photometry:** No training was necessary for fiber photometry studies, since the mice at no point were head restrained. The 5-day recovery period ensured that the circadian rhythm was restored, and that the surgical wounds were healed.

### DCE-MRI data collection

All MRI experiments were conducted on a 9.4 T preclinical scanner (Bruker BioSpec 94/30 USR, Bruker BioSpin, Ettlingen, Germany), interfaced with a Bruker Avance III console and controlled by ParaVision 6.0.1 software (Bruker)^12^. Imaging was performed using a room-temperature volumetric Tx/Rx resonator (35-mm inner diameter) and a 1,500 mT/m gradient coil (BFG6S, Bruker) in both the awake state and under ketamine/xylazine anesthesia (K/X: 100/10 mg/kg, intraperitoneally). For animals receiving K/X, an intraperitoneal catheter was implanted to allow supplementary delivery of half the dose of K/X every 50 minutes.

All animals were securely fixed to an MRI-compatible animal holder, with the implanted headplate positioned at a 30° neck angle in the prone position. The animal’s body was stabilized using a standard stretch cotton bandage. An MR-compatible remote monitoring system (SA Instruments, NY, USA) was employed to maintain body temperature at 37 ± 0.5°C using a thermostatically controlled waterbed, along with continuous monitoring of respiratory rate.

First, a morphological reference 3D constructive interference steady-state (3D-CISS; repetition time [TR]: 3 ms, echo time [TE]: 1.5 ms, number of averages [NA]: 1, flip angle [FA]: 50°, field of view [FOV]: 19.2 mm × 19.2 mm × 16 mm, matrix: 192 × 192 × 160, voxel size: 0.1 mm × 0.1 mm × 0.1 mm, maximum intensity projection from 4 TrueFISP volumes of orthogonal phase encoding directions) was acquired before the MR contrast agent injection. Secondly, the dynamic contrast-enhanced MRI (DCE-MRI) was performed using the protocol adapted from previous work ^20^, to produce quantitative and dynamic measurement of gadolinium concentration in brain tissue after intrastriate injection of the contrast agent. To correct for inhomogeneous RF pulse propagation, an RF transmit-field inhomogeneity map (B1^-^map) was obtained using a double-angle method with a 2D rapid acquisition with relaxation enhancement turbo spin-echo sequence (2D-RARE; TR: 8000 ms, TE: 23 ms, NA: 1, RARE factor: 8, 45 slices, matrix: 96 × 96, in-plane resolution: 0.2 mm × 0.2 mm, slice thickness: 0.4 mm, FAs: 70° and 140°). A pre-contrast 3D T1 map was acquired using a variable flip angle fast low angle shot method (VFA-FLASH; TR: 14.88 ms, TE: 1.41 ms, NA: 1, matrix: 96 × 96 × 45, voxel size: 0.2 mm × 0.2 mm × 0.4 mm, FAs: 2°, 3°, 5°, 8°, 15°, 25°, and 50°, scan time per FA: 1.5 min). Afterwards, three baseline scans followed by 60 contrast-enhanced scans over a 90-minute follow-up session were acquired using the FLASH sequence with a single FA=15°. The T1-enhancing contrast agent gadobutrol (20 mM; Gadovist, Bayer Pharma AG, Leverkusen, Germany) was infused into the striatum (0.1 µL/min for 10 min, total infusion volume = 1 µL). For reference imaging, a phantom filled with 50 µM gadobutrol was placed in the FOV to allow T1 value normalization between each scan. Reference Tx power and receiver gain were kept constant during the scanning session, as determined during the first RARE acquisition.

### MR image processing and analysis

The MRI series for 3D-CISS volumes acquired with four orthogonal phase encoding directions and the 3D-FLASH volumes were corrected for motion and image bias field using Advanced Normalization Tools (ANTs) ^21^. The processed 3D-CISS images were computed as a maximum intensity projection. Subsequently, VFA-FLASH volumes and post-contrast FLASH volumes were coregistered to the 3D-CISS volumes. To suppress the noise in each time series, the Marcenko-Pastur PCA (MP-PCA) algorithm denoising has been used. The 3D brain T1 maps for pre-contrast injection were calculated using a set of seven flip angles in VFA-FLASH with myRelax ^22^ and in-house python scripts. Dynamic 3D gadolinium concentration maps were constructed using the pre-contrast T1 calculated from VFA-FLASH and the post-enhancement ratio calculated from the successive acquisition of FLASH at the single flip angle. Gadolinium concentrations were simulated from a phantom study using the same imaging parameters as the FLASH protocol in this study, with the relationship between concentration and r1 (1/T1) set as 3.0 mM/second. To measure Gd concentration over time, a seed coordinate was determined at the tip of the intrastriatal cannula in the image volume. From this point, four spherical regions of interest (ROIs) were created with increasing radii (0-2.0 mm in increments of 0.5 mm). To focus on the signals in the brain interstitium, CSF regions (ventricles and subarachnoid spaces) were excluded from each ROI, and the mean concentration was measured. ROI placement and concentration measurements were performed using Imalytics Preclinical software (ver. 3.1.1.4, Gremse-IT GmbH, Aachen, Germany). Results were considered significant for p<0.05 using non-parametric Mann-Whitney-U test.

### SPECT/CT data collection

Previously, we developed an *in vivo* glymphatic imaging platform utilizing dynamic SPECT/CT imaging to monitor the distribution of radiolabeled tracers in rats ^23^ that here were adapted to awake, sleeping and anesthetized mice ^12^.

After successful habituation, each mouse was placed into a custom-made paper tunnel, and its headplate was secured to a custom-built fixation system within the imaging holder. This combined use of the tunnel and headplate fixation served as a complementary restraint method, effectively preventing head movement and excessive body motion during SPECT imaging. As for MRI, the fixation system allowed for a 35-degree flexion of the head to achieve a position closer to the natural head posture of the animal.

Imaging was performed using the VECTor4CT system (MILabs, Utrecht, Netherlands), equipped with a high-energy ultra-high-resolution multi-pinhole collimator for mice (HE-UHR-M, 0.7 mm pinhole diameter) allowing for higher resolution imaging with sustained high sensitivity. Each imaging session began with an initial whole-body CT for anatomical reference, followed by a SPECT acquisition. During the first experimental phase, the SPECT radiotracer infusion commenced immediately after the animal was positioned, and a rapid planar X-ray whole-body localizer was obtained in both axial and sagittal views. A 50 µL Hamilton microsyringe (Sigma Aldrich) mounted on a micro-infusion pump was used to deliver 1 µL of the tracer ^99m^Tc-labeled diethylenetriaminepentaacetic acid (12.5 mg/ml, TechneScan DTPA, Curium Pharma; 489 Da) into the striatum at rate of 0.1 µl/min, via a PE-10 tube connected to the striatal cannula. The concentration activities of the tracer at the time of infusion varied (mean ± standard deviation: 1.073 ± 0.156 MBq/ml) and did not differ significantly among the animals (p=0.2009 between the 3 groups, using Kruskal-Wallis one-way ANOVA). Six minutes before the end of the infusion, the reference CT volume acquisition began (50 kV, 4.8 mAs, fast scan mode, continuous source-detector rotation, angle speed = 10°/sec, single bed position, Al500 filter, binning x22, 100 µm³ reconstructed resolution, 40 sec acquisition duration). Two minutes before the infusion ended, whole-body SPECT imaging commenced in list-mode (0-1200 keV window) and consisted of two consecutive frames, each lasting 7.5 minutes. The acquisition in two frames accounted for the tracer activities reaching the brain parenchyma with latency. Each SPECT frame was acquired in 50 bed positions, covering the field of view (FOV) from the prior CT scan.

The second experimental phase repeated the imaging protocol from the first, excluding the tracer delivery. The same FOVs were used for the CT and SPECT acquisitions in both the first and second scans, with adjustments made to account for FOV translation depending on the animal’s positioning in the holder. After the successful acquisition of the first SPECT/CT volumes, the PE-10 tube containing the remaining tracer was cut and sealed 1-1.5 cm above the external tip of the cannula, and the animals were returned to their home cage under continuous video monitoring. Mice in the K/X group (K/X: 90/10 mg/kg, i.p.) were kept at 37°C and were supplemented every 45 minutes with half the dose of K/X.

Reflexes, presence of respiration pattern and its depth, and the color of mucus membranes were monitored throughout the recordings to ensure the animals’ well-being.

### SPECT image reconstruction and analysis

All SPECT images were reconstructed using a dedicated MILabs software (ver. 12.55-st) using ⟨125.1|152.9⟩ keV the photopeak window with the center at 140 keV, and the background in the ranges of ⟨119.5 |125.1)and (152.9|158.5⟩keV. The images were reconstructed using a Similarity-Regulated Ordered Subsets Estimation Maximization (SROSEM) with an isometric voxel size of (200µm)^3^ and using 20 iterations, after which a scatter and CT volume-based non-uniform attenuation correction was performed. Subsequently the reconstructed SPECT volumes were spatially co-registered (MILabs software, rigid-body registration) to their parental CT volume from the corresponding experimental phase to match the spatial FOV extents and resolution of (100µm)^3^. To account for the effects of the awake animal movement during SPECT acquisition, the reconstruction was performed considering the activities from two frames jointly (i.e. 1 volume of 15 min). To account for potential latency in tracer delivery, the reconstructed CT and SPECT volumes from the first experimental phase were only used to confirm the placement of the intrastriatal cannula with insert within the brain parenchyma, and to confirm presence of the activity in the insert and the brain after the infusion. The analysis was based on the second phase experimental SPECT volume and this data was included in the analysis only if the animal underwent successful delivery of the tracer, as confirmed by visible activity in its SPECT image from the first experimental phase.

A total of 17 mice were included in the SPECT study. Two animals were excluded due to a lack of tracer activity in the cannula or brain parenchyma during the first phase scan. The reconstructed SPECT images from the second phase were decay-corrected based on the exact time difference between the start of the first and second phase SPECT scans. All SPECT images were smoothed in three dimensions using a 1 mm FWHM Gaussian filter kernel. In all SPECT images, tracer activities within the cannula were removed based on corresponding CT images. The background and cannula regions were semiautomatically defined using Imalytics Preclinical software (ver. 3.1, Gremse-IT GmbH, Germany) and further dilated by 2 voxels in each orthogonal direction to account for slight animal movements. The total ^99m^Tc activities present in the second phase SPECT images were measured within the brain parenchyma, considering only the brain parenchyma volume, which was manually defined in a representative CT template volume using ITK-SNAP **(Fig. 2c)** ^24^. The template volume was co-registered to the skull image from the CT scan of each animal separately, using an affine transformation. The volume of the activities present in the brain parenchyma in each animal were calculated as a total number of the second phase SPECT image voxels having non-zero intensity value, multiplied by the voxel resolution.

Similarly to that in MRI evaluation, a spread of the tracer from the infusion point (i.e., cannula tip) was measured separately in each animal by calculating the total sum of activities present in the spherical distance of 0-0.5 mm (0.53 mm^3^ volume), 0.5-1 mm (3.7 mm^3^ volume), 1-1.5 mm (10 mm^3^ volume), and 1.5-2 mm (19.4 mm^3^ volume) from the infusion point. The spherical volumes were defined manually in each case, by placing the same ’ball-shaped set of volumes-of-interest centered at the cannula tip visible in the second phase CT image. To verify existence of the tracer spread brain-wide, an additional 0.53 mm^3^ volume of interest was placed at the contralateral hemisphere. The total derived tissue activity volumes, and the spherically measured activities were compared between the awake, sleep, and K/X groups using non-parametric Kruskal-Wallis ANOVA with Dunn’s correction and two-way ANOVA with Bonferroni’s post-hoc. Statistical differences were considered for p<0.05.

### Fiber photometry data collection

Mice allocated to the awake and sleep groups were habituated for 24 hours to the cage and experimental room. One hour before the start of the experiment, the cannula of each mouse was connected to 60 cm of PE10 tubing (PE10-Cl-500, Scandidact), which was attached to a 10 µl syringe (#1700, Hamilton) mounted on an infusion pump (Pump 11 Elite, Harvard Apparatus). The syringe and tubing were filled with mineral oil (Sigma) and 1.5 µl of 10% (w/v) 3kD FITC dextran (D3306, Life Technologies), diluted in aCSF (149 mM NaCl, 3.5 mM KCl, 1.2 mM CaCl₂, 0.85 mM MgCl₂, pH = 7.2, 289 mOsm). An optical cable (Ø200 µm core, 0.22 NA FC/PC to Ø1.25 mm ferrule patch cable, Thorlabs), connected to a fiber photometry system (RZ10-X Lux-I/O Processor with excitation LEDs (405, 465, and 560 nm) and LUX photosensors (LxPS1), TDT), was attached to the mouse’s optical fiber. A camera (Brio 100 FullHD, Logitech) was positioned above the cages for motility analysis.

The anesthetized group received K/X anesthesia and was placed on a heating pad (37.2°C) to maintain body temperature. These mice were connected to the same pump and photometry equipment as the other groups. Every 10 minutes, the toe pinch reflex was assessed, and additional K/X was administered as needed. Once the anesthetized mice showed no reflexes, and the awake and sleep groups were habituated for 1 hour, a 5-minute photometry recording was taken to determine baseline fluorescent signals in the absence of tracer. The pump, camera, and optical measurements were then simultaneously and remotely activated to inject 1 µl (0.1 µl/min) of tracer, while leaving the mice undisturbed.

After 3 hrs of recording, the mice were decapitated, and their brains were collected and immersed in 4% paraformaldehyde (v/v in PBS). Blood samples (20 µl) were collected and diluted in 20 µl of heparinized saline. The trigeminal nerve and superficial lymph nodes were immediately imaged *in situ*, while the optic nerves were dissected out prior to imaging using a fluorescent macroscope (Leica M205 FA fluorescence stereomicroscope with an Xcite 200DC light source and A12801-01 W-View GEMINI (Hamamatsu)). The volume of aCSF containing FITC-dextran remaining in the tubing at the end of the experiment was measured to determine the actual volume and, thereby, the dose of FITC-dextran that was injected. The fluorescence signal of the supernatant from the 20 µl blood sample was measured using the CFX Connect™ Real-Time PCR Detection System (Bio-Rad). A subgroup of brains was processed for routine histology to evaluate the inflammatory response to the cannula implant.

To validate the signal measured from the SSS, a separate group of anesthetized mice was connected to an identical optic fiber placed over the SSS. Additionally, a second optic fiber (Ø400 µm core, 0.5 NA FC/PC to Ø2.50 mm ferrule patch cable, Thorlabs) was positioned against the sphincter of the rectum. Increasing concentrations (1%, 2%, and 4%) of 3kD FITC, diluted in 100 µl saline solution, were injected IV retro-orbitally at a volume of 1 µl. A second group of mice received an intrastriatal injection of 2 µl of 10% (w/v) 3kD FITC dextran (0.1 µl/min).

### Fiber photometry data analysis

The FITC-dextran signal over the superior sagittal sinus was calculated using the 465 nm signal recording, with movement artifacts removed based on the 405 nm signal recording and their respective baseline (MATLAB). The mean pixel intensity was calculated for the optic and trigeminal nerves, as well as the superficial lymph nodes, by segmenting the fluorescent tissue through thresholding (FIJI, ImageJ). All data were normalized to the awake group, with the mean of the awake group set to 1 as the reference point. Statistical significance was determined with p<0.05 using one-way ANOVA or two-way ANOVA with Bonferroni’s post-hoc test.

### Immunohistochemistry preparation and data analysis

Immunohistochemistry was performed as previously described ^13,25^. The primary antibodies used were rabbit anti–glial fibrillary acidic protein (GFAP; 1:250, Agilent Cat# Z0334, RRID:AB_10013382); goat anti–ionized calcium-binding adapter molecule 1 (Iba1; 1:500,Abcam Cat# ab5076, RRID:AB_2224402) and NeuN (goat, 1:500, Abcam Cat# ab177487, RRID:AB_2532109). GFAP, NeuN, and Iba1 immunofluorescence images were collected using a Nikon Eclipse Ti2 microscope. For image quantification, serial confocal images were acquired with the Pan Fluor ×40/1.30 NA objective (2 μm, 10 steps, excitation sources 405, 488, 565nm, and 647 solid-state diode laser lines).

The max projection of the entire z-stack on the area around the cannula implant (striatum, ipsilateral, IP) or the contralateral side (striatum, CL) were used to calculate % area covered by Iba1 or GFAP by using a fixed max entropy thresholding (ImageJ, NIH). Neuronal cell density was calculated based on a particle analysis of the NeuN+ cells. Each mouse produced a single biological replicate.

## Author contribution

EK, LVR, RSG, SAF, MN, PW, YM, VP, TE, VHS, EBK, ESN performed the experiments, collected the data, and performed the analysis. YM, SAG and MN wrote the manuscript. All authors read and approved the final version of the manuscript.

## Data and code availability

Requests for data from this study and further information should be directed to the corresponding author, M.N. All unique resources generated in this study are available from the corresponding author with a completed materials transfer agreement.

## Acknowledgement

We thank Dan Xue for excellent graphic support. We also thank Helle Hvorup Knudsen and Martin Dan Tramm W Havndrup for superb technical support. This project received funding from the Lundbeck Foundation (R386-2021-165 to M.N), NNF (NNF20OC0066419 to M.N.), the NIH (R01AT012312RF1AG057575, R01AT011439, U19 NS128613 to M.N.), the US Army under award MURI W911NF1910280 (to M.N.), the Simon and Adelson foundations (to M.N.), JPND/HBCI 1098-00030B (M.N.); JPND/Good Vibes 2092-00006B (M.N.), and from National Center for Complementary and Integrative Health grants (NCCIH R01AT011439 and R01AT012707 to M.N.).

